# DeepKOALA: A Fast and Accurate Deep Learning Framework for KEGG Orthology Assignment

**DOI:** 10.64898/2026.01.07.698072

**Authors:** Zhaoxi Yu, Lingjie Meng, Canh Hao Nguyen, Hiroshi Mamitsuka, Minoru Kanehisa, Hiroyuki Ogata

**Affiliations:** Bioinformatics Center, Institute for Chemical Research, Kyoto University, Gokasho, Uji 611-0011, Japan

**Author notes:** Correspondence should be addressed to Hiroyuki Ogata.

## Abstract

The KEGG Orthology (KO) system links DNA and protein sequences to biological functions and pathways, providing a curated, fundamental and consistent annotation framework across all domains of life. While accurate, traditional sequence alignment-based annotation methods are computationally expensive, which severely limits their application in large-scale datasets. To address this challenge, we introduce DeepKOALA, a deep learning approach based on Gated Recurrent Units (GRU), which frames KO annotation as an open-set recognition task. This design reduces false positives arising from out-of-scope sequences and, together with a lightweight GRU backbone, enables high-throughput annotation. The GRU-based model was benchmarked against four other deep learning architectures and showed the best balance between speed and accuracy. We further performed a cross-species evaluation of DeepKOALA in comparison with existing KO annotation tools, where it achieved an F1-score of 0.8653 and 37.5-fold acceleration compared with BlastKOALA. We also provide a specialized fragment model for handling incomplete sequences and an optional Multi-domain mode. Together, these features make DeepKOALA a scalable, efficient, and accurate solution for high-throughput function annotation.

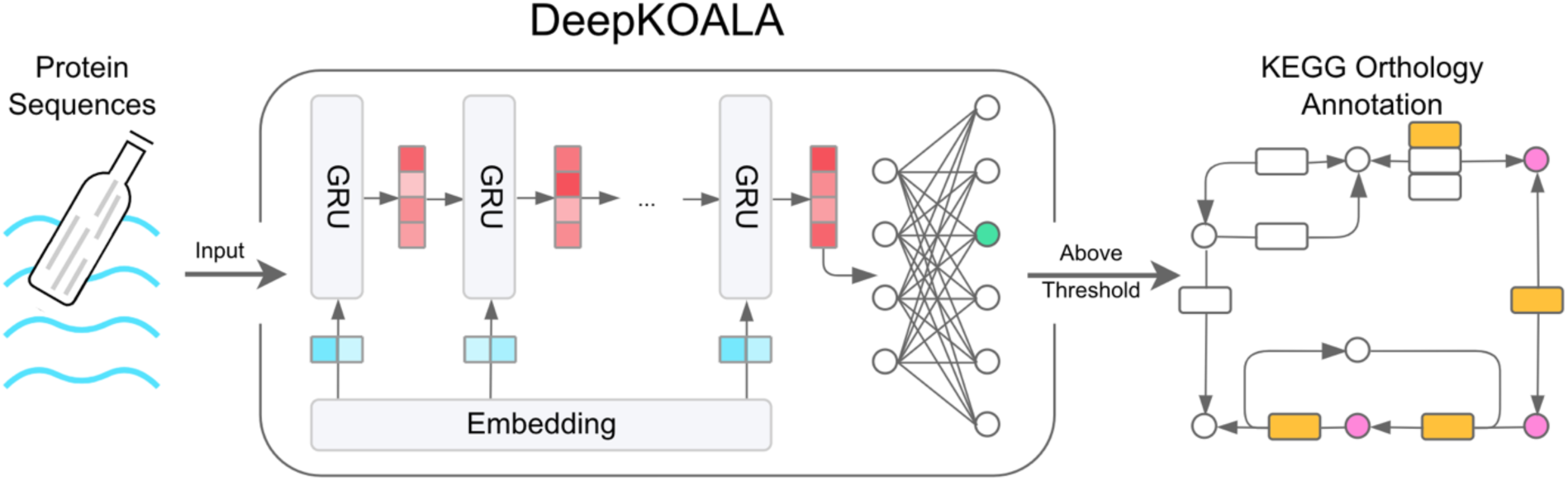

## Introduction

The Kyoto Encyclopedia of Genes and Genomes (KEGG) and KEGG Orthology (KO) system provide foundational resources for the systematic understanding of gene functions [1]. Built through manual curation, the KO system links genes to diverse biological data organized in an experiment-grounded comprehensive framework. Current KO annotation relies on two sequence-alignment-based approaches. The first detects homologous sequences via pairwise sequence alignment—e.g., BLAST-based workflows such as BlastKOALA and GhostKOALA [2] — whose runtime scales with the reference database size. The second approach scans sequences against KO-specific profile hidden Markov models (HMMs)—e.g., KofamKOALA/KofamScan [3]—whose runtime scales with the total number of KO profiles, thus reducing the computational cost. However, these alignment-based methods face a serious computational bottleneck for large-scale analyses. This challenge is particularly acute in metagenomics, which produces petabyte-scale datasets comprising vast sequence spaces of both “known” and “unknown” functions. There is therefore an urgent need for an annotation approach that is both efficient and accurate at scale.

Recent advances in computational biology have introduced a range of deep-learning-based architectures for protein sequence modeling and protein family classification, including Convolutional Neural Networks (CNN) [4], Recurrent Neural Networks (RNN) [5], Gated Recurrent Units (GRU) [6], Long Short-Term Memory (LSTM) [7], state-space models (e.g., Mamba [8]), and large protein language models (e.g., ESM-2 [9]). Many of these efforts have focused on Gene Ontology (GO) annotation, with representative methods such as DeepGO [10], DeepGOPlus [11], DeepFRI [12], TALE [13], ProLanGO [14], DeepGraphGO [15], NetGO [16], GOLabeler [17] and the zero-shot/neuro-symbolic approaches DeepGOZero [18] and DeepGO-SE [19]. Beyond GO, deep-learning-based models also annotate enzyme EC numbers (e.g., ProteInfer [20] and more recent trustworthy/uncertainty-aware frameworks) and antibiotic resistance genes (e.g., DeepARG [21], HMD-ARG [22]). Although KO remains one of the most widely used systems for pathway-level annotation, existing KO prediction tools rely on sequence alignment and could be improved by deep-learning solutions. Furthermore, most previous alignment-based and machine learning-based approaches formulate prediction as a closed-set problem, assuming that all query sequences belong to known classes, thereby lacking mechanisms for calibrated rejection of out-of-scope sequences [23,24].

We address this limitation by framing KO annotation as an open-set recognition problem and present DeepKOALA. To effectively reject sequences with unknown functions, we calibrate per-class, reliable decision thresholds by jointly leveraging labeled proteins and a large collection of unlabeled (unknown) sequences. We benchmarked RNN, GRU, LSTM, Mamba, and ESM-2 (8M; checkpoint: esm2_t6_8M_UR50D), and selected GRU as the backbone based on accuracy–efficiency trade-offs and broad CPU/GPU compatibility. DeepKOALA achieves accuracy comparable to existing alignment-based tools, while substantially improving throughput. DeepKOALA also includes two practical features for fragmented sequences and multi-domain proteins. We provide a user-friendly web server on GenomeNet (https://www.genome.jp/tools/deepkoala/), downloadable DeepKOALA software on GitHub (https://github.com/zhaoxi120/deepkoala), and regularly updated weights for the models.

## Materials and Methods

### Overall Workflow

In the model development phase, we benchmarked several architectures on KEGG sequences and selected a lightweight GRU for its optimal balance of accuracy and efficiency. We formulated KO assignment as an open-set recognition problem, where the goal is to correctly identify known functions while rejecting unknown ones using decision thresholds. To determine these thresholds effectively, we calibrated them for each KO class using both labeled and unlabeled sequences. During the prediction phase, the trained model encodes each input sequence into a fixed-length vector, which is mapped to KO probabilities by a multilayer perceptron (MLP). The top-scoring KO is assigned only if its probability exceeds the calibrated threshold; otherwise, the sequence is rejected as unknown.

### Dataset Construction and Preprocessing

Our study is framed within the KO system (February 2025), which defines 26,226 functional classes. We utilized the KEGG GENES database (February 2025) with a collection of 55,795,412 protein sequences, of which 52.35% (29,207,030 protein sequences) were annotated with KOs while others were unclassified (26,588,382 protein sequences). To mitigate extreme class imbalance, we applied a targeted augmentation strategy for underrepresented KO classes (see Supplementary Method S1 for details).

To mitigate the bias from sequence redundancy, we used the CD-HIT [25] tool (with parameter – c 0.9) to cluster the combined dataset and removed redundant sequences with over 90% identity. This resulted in a non-redundant dataset of 31,716,962 sequences (13,682,867 with KO labels and 18,034,095 without). For protein sequences associated with multiple KO labels, we assigned each sequence a single class label corresponding to the unique combination of its KOs (a total of 243 such combination classes). We employed a targeted oversampling strategy to address data imbalance within the training splits. Specifically, for any KO class (including the combination classes) that contained very few sequences in a given training split, we repeatedly and randomly duplicated its existing member sequences until the total count for that class in that split reached a phase-specific minimum (see Supplementary Method S1 for details).

### Model Optimization and Selection

To identify the optimal deep learning architecture, we systematically compared five candidate models: RNN, GRU, LSTM, Mamba, and ESM-2 (8M). The selection process involved two key stages: hyperparameter optimization and cross-validation. For this two-stage process, the full dataset was randomly divided into a 10% subset dedicated to hyperparameter optimization and a 90% main evaluation set reserved for the subsequent cross-validation.

We first used a dedicated hyperparameter tuning set and employed a Bayesian optimization algorithm to find the optimal configuration for each model. This optimization was guided by the Open-set Area Under the Curve (OpenAUC) [26], a threshold-independent metric specifically suited for open-set recognition problems such as the KO annotation task in this study. In this setting, a model must not only classify known KOs but also reject unknown sequences.

Mathematically, it is defined as:

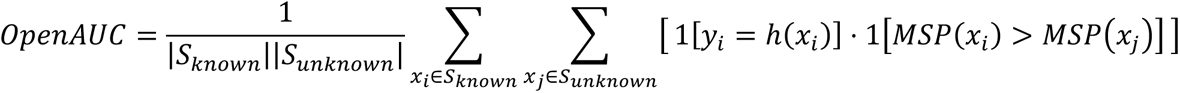

where *S_known_* and *S_unknown_* are the sequence set with and without KO labels, respectively. ℎ(*x_i_*) and *y_i_* are the predicted and true labels for *x_i_*. *MSP* is the maximum Softmax score. In this formula, the model’s performance is evaluated on a pair (*x_i_*, *x_j_*) of sequences with and without KO labels. When the model both assigns the correct label to the known sequence (i.e., *y_i_* = ℎ(*x_i_*)) and assigns a higher Softmax score to the known sequence compared to the unknown one (i.e., *MSP*(*x_i_*) > *MSP*(*x_j_*)), we consider this pair as the correct pair. The computed OpenAUC is the probability of correct pair over all known-unknown pairs. A higher OpenAUC score thus indicates a proper balance between classification accuracy and the ability to reject unknown sequences, making it the core metric for our performance evaluation.

To strike a balance between model performance (OpenAUC) and computational efficiency (time), we defined a selection criterion function *f*(*OpenAUC*, *Time*) to identify the optimal configuration. For each hyperparameter combination, we first performed a min-max normalization on the OpenAUC scores and time consumption:

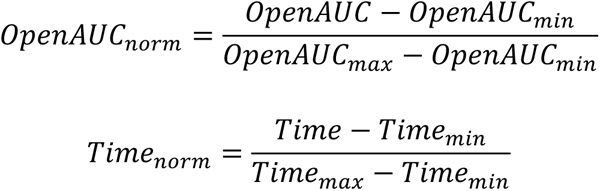

We then calculated the cost value *f*(*OpenAUC*, *Time*) for each configuration, representing its deviation from the ideal scenario (where *OpenAUC* = 1 and *Time* = 0). Reflecting the principle that annotation accuracy is the primary consideration in functional genomics, we assigned a performance weight (*W_openAUC_*) of 0.8 and a time weight (*W_Time_*) of 0.2. The criterion function is defined as:

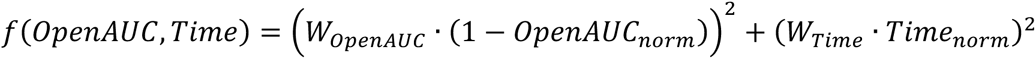

Finally, the configuration that minimized the cost function *f*(*OpenAUC*, *Time*) was selected as the optimal setting for each model. Ideally, the best configuration for each of the four most promising models (RNN, GRU, LSTM, Mamba) was used for a rigorous five-fold cross-validation on the main evaluation set. The final model architecture was chosen based on the best balance of OpenAUC performance, computational efficiency, and versatility across different hardware environments (CPU/GPU).

### Model Framework and Algorithm

The model selected in this study is a GRU-based deep learning network where the detailed structure is as follows (Figure 1B):

● **Input Layer:** Accepts variable-length protein sequences.
● **Embedding Layer:** Maps each amino acid residue to a learned high-dimensional vector to capture its physicochemical properties.
● **GRU Layers:** Two stacked GRU layers. The first layer encodes residue context from left to right. Its full output sequence is fed to the second layer to learn higher-level features. For classification, we take the last hidden state as the sequence embedding.
● **Output Layer:** The embedding is fed into a one-hidden layer MLP, which maps the sequence embeddings to KO classes with raw output scores.
● **Prediction Layer:** Finally, a Softmax activation function converts the raw output scores into predicted probabilities for each KO class.

**Figure 1.**
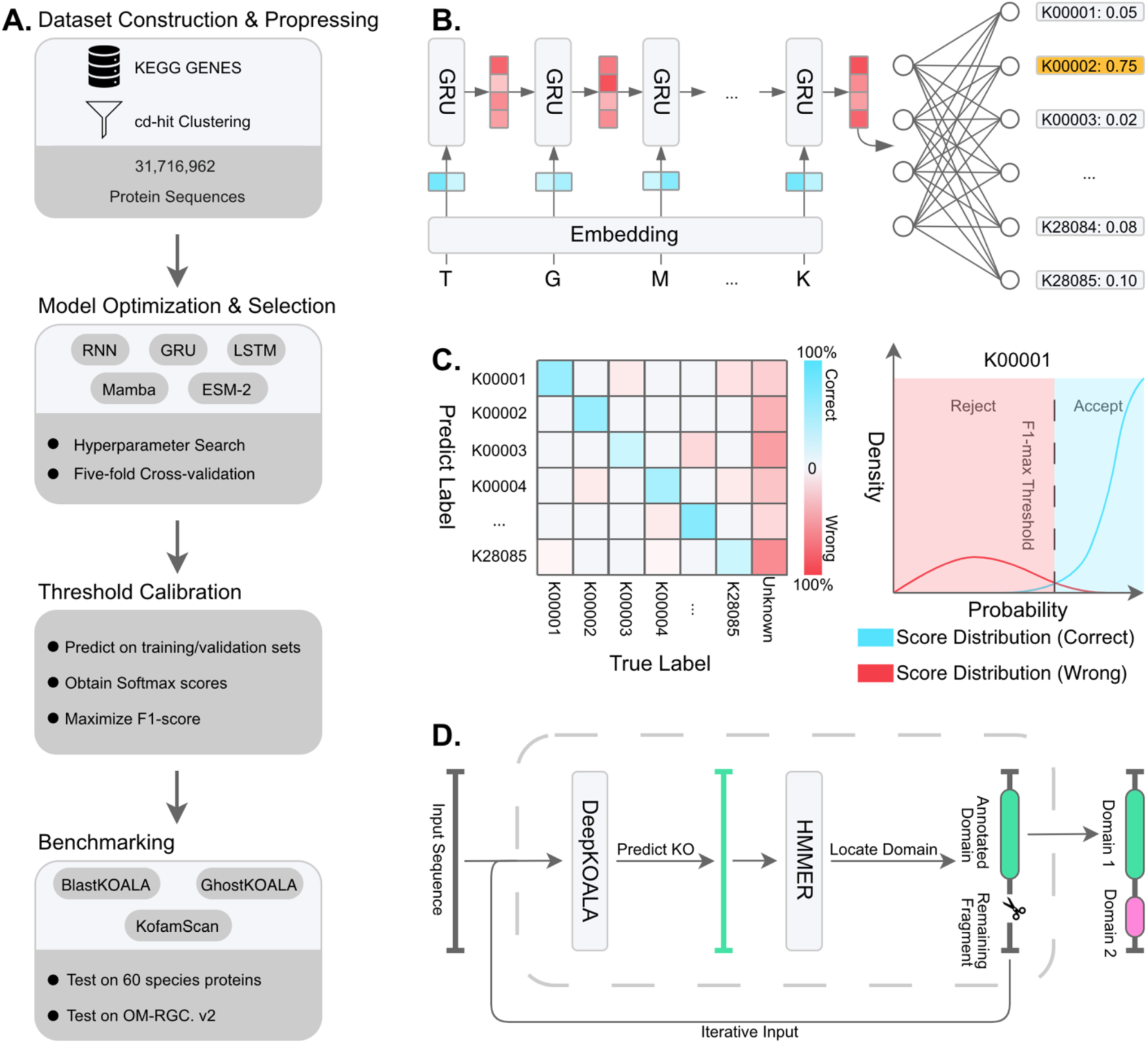
Overview of the DeepKOALA framework. **A**, The workflow pipeline comprises four main stages: dataset construction & preprocessing, model optimization & selection via hyperparameter optimization, threshold calibration, and final benchmarking against existing annotation tools. **B,** The model architecture utilizes an embedding layer and two stacked GRU layers to encode protein sequences, which are then processed by a MLP and Softmax to output KO class probabilities. **C,** Threshold determination mechanism: the confusion matrix (left) categorizes predictions into correct (blue) and wrong (red) groups, while the density plot (right) analyzes their score distributions to identify the optimal threshold maximizing the F1-score. **D,** The multi-domain mode employing an iterative hybrid strategy, where DeepKOALA predicts the KO label and HMMER delineates the domain boundary, allowing the annotated domain to be computationally removed and the remaining fragment to be re-analyzed for additional domains.

### Threshold Calibration

To retrain the selected GRU model, the full dataset was randomly divided into three sets: training (70%), validation (10%), and test (20%) set. While the threshold-independent OpenAUC metric is ideal for model comparison, converting the final model into a practical annotation tool requires establishing a specific decision threshold for each KO class. This threshold is applied to the output Softmax probability to define positive (“known”) and negative (“unknown”) predictions.

To determine the optimal threshold for each KO class, we implemented a systematic procedure. First, we used the trained model to generate predictions and corresponding Softmax scores for all data in the training and validation sets. For each KO class, we focused on the sequences predicted to belong to that class and divided them into two groups: those whose true labels matched the predicted KO class, and those whose true labels differed from it (including sequences annotated with other KO classes or unlabeled sequences). Based on the Softmax score distributions of these two groups, we determined the final threshold score that maximized the F1-score (the harmonic mean of precision and recall) for the KO class (Figure 1C, Supplementary Method S4).

### Final Performance Benchmarking

After selecting the optimal GRU-based architecture, we retrained the model on the full dataset and benchmarked its performance against three KO annotation tools: BlastKOALA, GhostKOALA, and KofamScan. To ensure a rigorous and fair comparison, we generated an independent test gene set of 60 species (N = 454,580 sequences), including prokaryotes and eukaryotes. The list of test species and excluded reference species are summarized in Supplementary Tables S3 and S4. All sequences from these species and their genera were strictly removed from the training and validation dataset and the reference databases of all tools prior to evaluation, to prevent data leakage and assess true generalization performance (see Supplementary Method S2).

### Additional Features

To enhance the model’s practical applicability, we introduced two auxiliary features:

● **Fragment model (DeepKOALA-fragment):** An independently trained variant model using cropped sequences to improve robustness to incomplete sequences (see Supplementary Method S3).
● **Multi-domain mode:** This mode employs an iterative hybrid strategy, combining the rapid deep learning model with the precise scanning of HMMER [27], to achieve comprehensive annotation of multiple functional domains on a single protein (Figure 1D; see Supplementary Method S3 for the detailed pipeline).

## Results

### Model Architecture and Hyperparameters Selection

To identify the optimal configuration that balances performance and efficiency for four candidate deep learning models (RNN, GRU, LSTM, and Mamba), we conducted a systematic Bayesian optimization search. Key hyperparameters, including the number of stacked layers, hidden units, and the learning rate, were tuned for each model. As illustrated in the performance-efficiency trade-offs (Figure 2), both GRU and LSTM models achieved the highest OpenAUC scores (∼0.91) under optimal settings, with the GRU models requiring less execution time. The Mamba model’s performance was close behind with an OpenAUC of 0.90, while the RNN model was lower at approximately 0.86.

**Figure 2:**
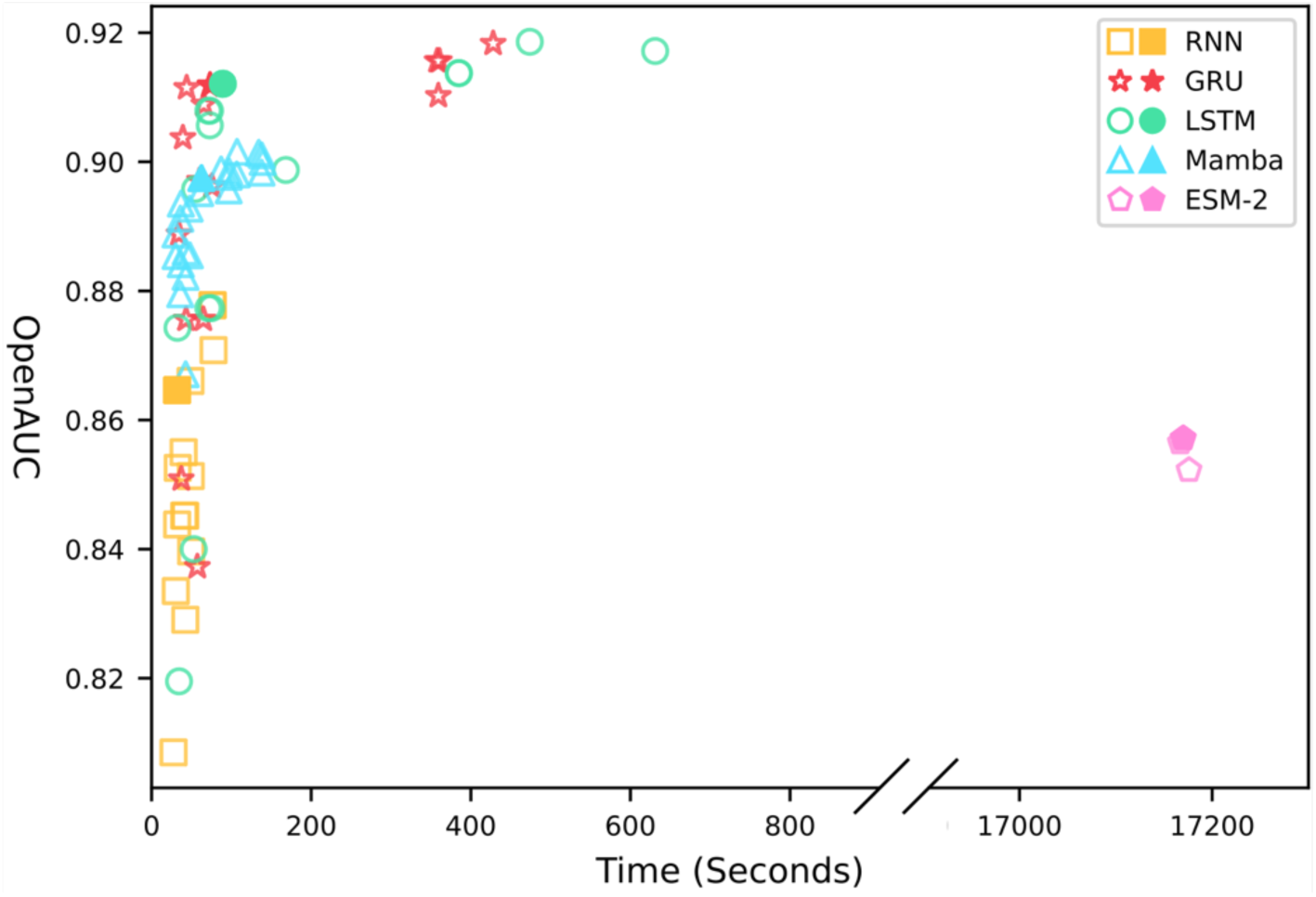
Performance-efficiency distribution from the hyperparameter search. The plot shows the performance of the five models under different hyperparameter combinations. Each data point represents a hyperparameter configuration, with its y-coordinate as OpenAUC and x-coordinate as time cost (seconds). The x-axis has a break to show the substantial time cost of ESM-2 (8M). Solid data points indicate the selected optimal configuration for each model.

Separately, we evaluated the pre-trained ESM-2 (8M) model, restricting optimization to tuning the learning rate of the final MLP classifier rather than fine-tuning the entire model. Despite its computation time far exceeding the other models up to 100-fold, ESM-2 (8M) yielded no performance advantage, achieving an OpenAUC score of around 0.86. Consequently, ESM-2 (8M) was excluded from subsequent cross-validation. Based on these results, we selected the optimal hyperparameter combination for each of the four remaining models (Table 1) to proceed to the next stage of evaluation.

**Table 1:**
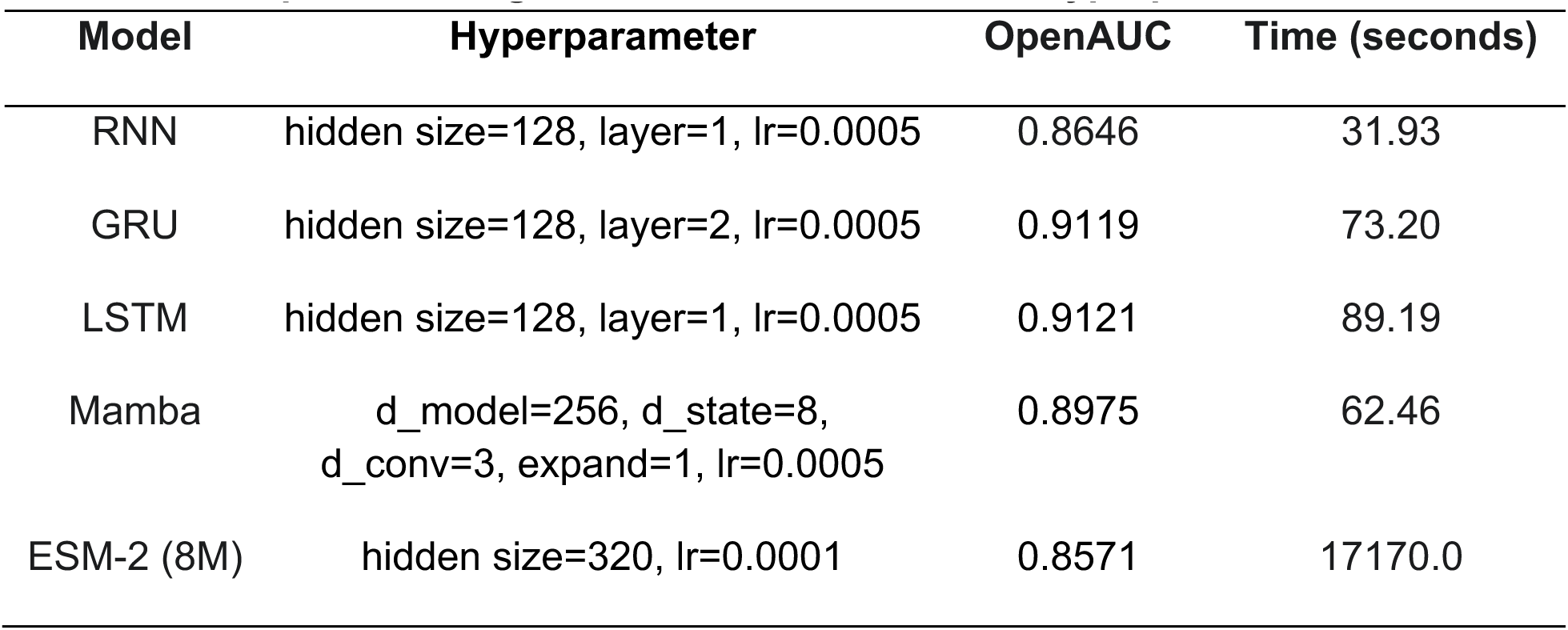
Optimal configurations selected from the hyperparameter search.

### Model Selection using Cross-Validation

After identifying optimal hyperparameters, the four most promising models were subjected to a rigorous five-fold cross-validation. As shown in Table 2, GRU and Mamba outperformed RNN and LSTM. Although GRU was ∼16% slower than Mamba (328.06 s vs 282.49 s), GRU achieved higher OpenAUC (0.9493 vs 0.9469). In addition, the GRU model also demonstrated better versatility by running efficiently on both CPU and GPU environments, whereas Mamba can run only on GPU. Given the strong performance, broader hardware compatibility, and suitability for deployment in resource-constrained settings, the GRU model was selected as the best solution for KO annotation.

**Table 2:** Five-fold cross-validation results for selected deep learning models.

| Model | OpenAUC | Accuracy | AUROC | Time(s) |
| --- | --- | --- | --- | --- |
| RNN | 0.8867±0.0005 | 0.5717±0.0185 | 0.7236±0.0050 | 140.09±1.81 |
| GRU | 0.9493±0.0002 | 0.9445±0.0004 | 0.9271±0.0004 | 328.06±1.01 |
| LSTM | 0.9421±0.0009 | 0.9154±0.0009 | 0.9085±0.0015 | 399.25±3.40 |
| Mamba | 0.9469±0.0007 | 0.9451±0.0007 | 0.9234±0.0008 | 282.49±1.40 |

### Threshold Calibration for KOs

To obtain the final prediction model, we first retrained the GRU network on the full dataset. When evaluated on the test set in a threshold-independent manner, the model achieved a high OpenAUC of 0.9495, an accuracy of 0.9446 on labeled sequences, and an AUROC of 0.9274 for discriminating between labeled and unlabeled sequences. We mainly compare methods using two metrics, F1-score and specificity: F1-score evaluates KO assignment performance on labeled sequences, whereas specificity evaluates unknown rejection as the fraction of unlabeled (unknown) sequences correctly rejected (not assigned to any KO), helping control false-positive KO assignments on out-of-scope sequences.

To convert these probability scores into practical class assignments, we systematically evaluated three distinct thresholding strategies: treating all unlabeled sequences as a single “unknown” class, the OpenMax framework [24], and our proposed joint labeled–unlabeled calibration.

We first examined the two common open-set approaches. The strategy of treating all unknown sequences as a single class achieved high specificity (0.9535) but failed to effectively recall known KOs, resulting in the lowest F1-score (0.8194). Conversely, the OpenMax method (p=5), which estimates thresholds based solely on labeled data distribution, suffered from the opposite problem. While it yielded a relatively high F1-score (0.8934), it suffered from poor specificity (0.7567), indicating a high rate of false-positive annotations.

To address these limitations, we implemented the joint labeled–unlabeled calibration strategy. Unlike the previous methods, this approach calibrates per-class thresholds by explicitly modeling the score distributions of both labeled (known KO) and unlabeled (unknown) sequences. Within this framework, we tested optimizing for accuracy, Youden’s index, and F1-score. As shown in Table 3, optimizing for the F1-score achieved the most favorable trade-off (F1 = 0.8653; Specificity = 0.9049). In contrast, optimizing for accuracy skewed the model toward negative predictions, increasing specificity to 0.9319 but dropping the F1-score to 0.8420.

**Table 3:** Performance comparison under different threshold-calibration strategies.

| Thresholding Metric | F1-score | Specificity |
| --- | --- | --- |
| F1-optimized strategy | 0.8653 | 0.9049 |
| Accuracy-optimized strategy | 0.8420 | 0.9319 |
| Youden’s Index-optimized strategy | 0.8589 | 0.8769 |
| Unknown as a Single Class | 0.8194 | 0.9535 |
| OpenMax (p=5) | 0.8934 | 0.7567 |
| OpenMax (p=10) | 0.8436 | 0.8422 |

The results demonstrate that approaches relying solely on labeled data or aggregating all unknown sequences into a single class yield less balanced decision boundaries. By jointly leveraging labeled and unlabeled data, our F1-optimized strategy effectively regularizes the thresholds, establishing reliable decision boundaries.

### Benchmarking Against Homology-based Tools

To evaluate the practical performance of our final model, we benchmarked DeepKOALA against three existing annotation tools—BlastKOALA, GhostKOALA, and KofamScan in a CPU environment (see Supplementary Method S2 for hardware, software, and benchmarking details). While BlastKOALA achieved the highest overall precision, DeepKOALA demonstrated a precision comparable to the alignment-based tool GhostKOALA (Figure 3). Specifically, DeepKOALA correctly annotated 143,975 sequences—a number nearly identical to KofamScan’s 145,459—but did so with a significantly higher precision (DeepKOALA 84.13% vs. KofamScan 78.74%), indicating fewer false positives. This high-precision performance is coupled with exceptional computational efficiency: DeepKOALA was 37.5 times faster than BlastKOALA, 6.7 times faster than GhostKOALA, and 6.3 times faster than KofamScan. On a single NVIDIA H100 GPU, the same dataset can be annotated in 66 seconds, providing an additional order-of-magnitude acceleration without altering the prediction outputs. This balance of high precision and speed underscores the practical advantages of our deep learning approach. Length-dependent accuracy profiles on the 60-species test set are shown in Supplementary Figure S3.

**Figure 3:**
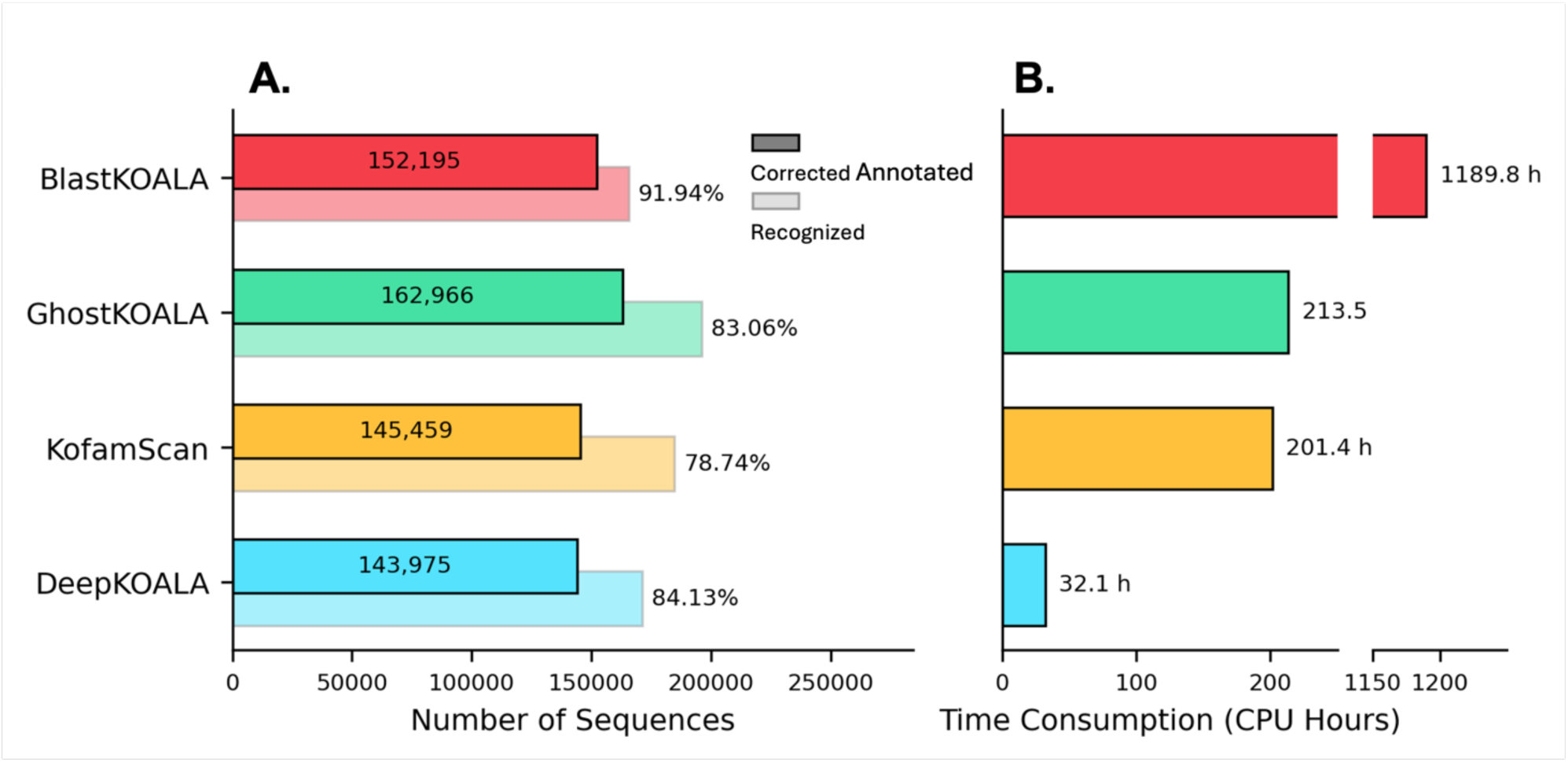
Performance and efficiency comparison with existing tools. **A**, Comparison of the number of sequences recognized (light bars) and correctly annotated (dark bars) by DeepKOALA, KofamScan, GhostKOALA, and BlastKOALA on the independent test set of 60 species. **B,** The corresponding runtime comparison of these tools, highlighting the computational efficiency of DeepKOALA compared to alignment-based methods.

### Application in Metagenomic Datasets

To assess DeepKOALA’s performance on complex, large-scale metagenomic data, we first tackled the issue of sequence fragmentation, a common problem in real metagenomic data caused by ORF misannotation or gene degeneration. We compared our standard DeepKOALA with a specialized variant model, DeepKOALA-fragment, that was trained on protein sequences cropped to variable lengths. On a test set of full-length proteins, both models showed comparable performance, with the standard DeepKOALA achieving a slightly higher accuracy (86.52% vs. 84.99%) (Figure 4A). However, when we evaluated models on a fragmentation-simulation test set using randomly cropped sequences, the DeepKOALA-fragment model outperformed the standard model. It not only achieved higher accuracy (86.07% vs. 83.85%) but also recognized substantially more sequences (1,948,803 vs. 1,481,137) (Figure 4B).

**Figure 4:**
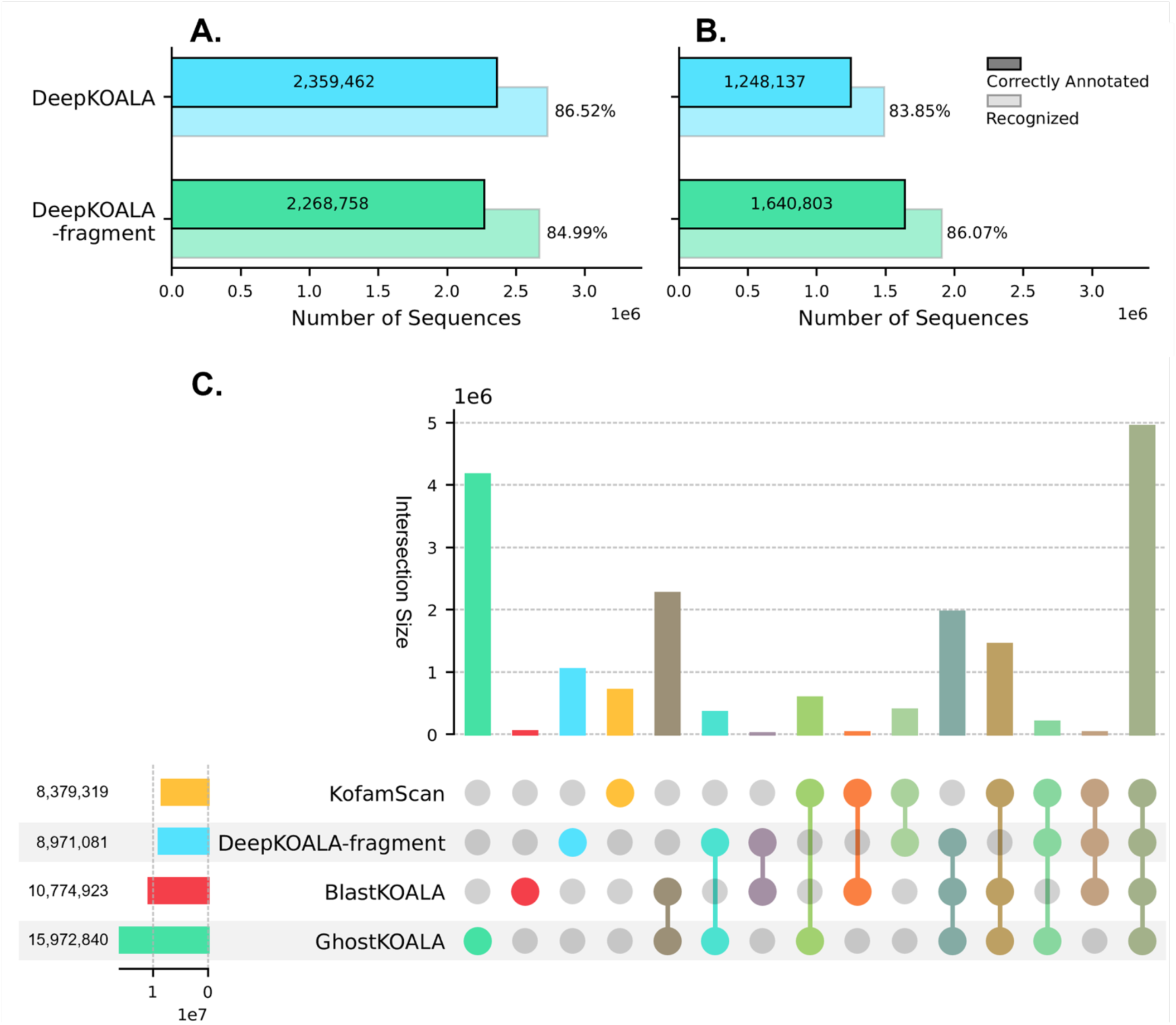
Performance of DeepKOALA on Metagenomic Datasets. **A**, Performance comparison between the standard DeepKOALA and the DeepKOALA-fragment model on a full-length protein test set, showing comparable accuracy. **B,** Performance comparison on a fragmented protein test set, demonstrating the superior robustness of the fragment model. **C,** An UpSet plot showing the intersection of annotations by DeepKOALA-fragment, KofamScan, BlastKOALA, and GhostKOALA on the OM-RGC.v2 dataset; the bars represent intersection sizes, with the single dot column indicating annotations unique to DeepKOALA.

Building on this, we applied the DeepKOALA-fragment model to the entire Ocean Microbial Gene Catalog (OM-RGC.v2) [28], which contains 46,775,154 protein sequences. We compared the annotation coverage of DeepKOALA-fragment with KofamScan, BlastKOALA, and GhostKOALA (Figure 4C). The analysis revealed a substantial core set of 4,950,790 sequences co-annotated by all four tools, confirming that DeepKOALA reliably identifies conserved proteins. Additionally, 1,041,855 sequences were exclusively annotated by DeepKOALA but missed by the other three methods, highlighting its potential in uncovering novel functional information within complex datasets. In terms of runtime, DeepKOALA demonstrated exceptional scalability, completing the task in just 38 minutes and 35 seconds on a single NVIDIA H100 GPU. For comparison, the same job would require an estimated 1,172 CPU hours on a CPU-only system for DeepKOALA, representing a 1,850× speedup.

### Multi-domain Mode for Multi-domain Proteins

Both the standard DeepKOALA model and the fragment model assign a single KO label for each input sequence, which limits their ability to identify multi-domain proteins. To overcome this, we developed a Multi-domain mode that combines the GRU model with HMMER in an iterative search strategy, in order to identify multiple domains within a single sequence (Figure 1D).

We validated this mode using a dataset of 152 RNA polymerase sequences, a family known to include both single– and multi-domain members. As an example, an RNA polymerase (1277 amino acid residues) from *Caldivirga maquilingensis* containing both K03041 (rpoA1; DNA-directed RNA polymerase subunit A’) and K03042 (rpoA2; DNA-directed RNA polymerase subunit A”) domains was processed using this mode. In this iterative procedure, DeepKOALA first assigned the KO label K03041, and HMMER was then used to delineate the corresponding domain boundaries (residues 15–902). The K03041 segment was subsequently removed computationally before the next iteration. Subsequently, the remaining sequence was re-analyzed and the K03042 domain was correctly identified (residues 910-1275). The consistent success on both single– and multi-domain sequences in our test set confirms the efficacy of this iterative strategy (Supplementary Table S9). It effectively resolves the false negatives inherent in single-label classifiers, enabling more comprehensive and accurate functional annotations for complex proteins. To evaluate the computational cost, we applied multi-domain mode to the 60-species independent test set. The entire dataset was processed in 76.7 CPU hours.

## Discussion

Protein annotation at scale remains a major bottleneck in large metagenomic data analysis, where existing tools struggle with speed, accuracy, or scalability. In this study, we presented DeepKOALA, a deep learning-based tool for KEGG Orthology annotation. By framing KO assignment as an open-set recognition problem, DeepKOALA achieves strong performance on a cross-species benchmark and delivers higher precision than KofamScan (84.13% vs 78.74%) at a similar level of true positive predictions, while it was comparable to GhostKOALA in terms of precision. Although BlastKOALA attained the highest precision (91.94%), DeepKOALA was 37.5 times faster than BlastKOALA on CPU. DeepKOALA annotated the entire OM-RGC.v2 database, containing about 47 million sequences, in approximately 38 minutes on a single GPU or 1172 CPU hours on a CPU-only system. Thus, DeepKOALA couples high accuracy with calibrated open-set rejection and high throughput for large genomic and metagenomic datasets.

The advantages of DeepKOALA stem from a fundamental shift in its computational paradigm and careful model architecture selection. DeepKOALA’s inference time is solely dependent on the sequence length and independent of the total number of KO classes, ensuring excellent scalability. Regarding model selection, we considered both large language models (e.g., ESM-2) and lightweight recurrent networks (e.g., GRU). Large models learn rich, context-aware, general-purpose representations from massive unlabeled data via pre-training, making them powerful for feature extraction tasks like protein structure prediction. However, for a supervised classification task with tens of thousands of fine-grained labels in our study, the goal is to learn highly specific, discriminative features. In this context, the massive size and quadratic complexity (*O*(*L*^2^)) of ESM-2 result in substantial time costs without a corresponding gain in classification performance. Consequently, a lightweight GRU backbone—with linear complexity (*O*(*L*)) and a small parameter footprint—provides the best accuracy–throughput trade-off for sequence-to-label assignment in our setting.

On a deeper level, KO classification presents challenges that are particularly well-suited to deep learning. The KEGG Orthology system is itself a manually curated, hierarchical classification system designed to represent biological function. This means the degree of internal sequence similarity greatly varies among different KO families, and simple sequence similarity is not always a reliable basis for classification. Deep learning models, on the other hand, can adaptively learn and adjust optimal recognition thresholds based on the underlying distribution of each KO class. This allows them to define the boundaries across KO families beyond simple sequence conservation. To address this variability, we calibrate per-KO thresholds in a semi-supervised manner: labeled sequences locate the F1-optimal operating point, while unlabeled sequences help regularize against false positives. Compared with a fixed global threshold or OpenMax-style thresholds estimated from labeled data alone, this yields a more stable precision–recall/specificity trade-off. It also aligns with the practice of class-specific adaptive thresholds in KO pipelines.

There are several inherent limitations in the current DeepKOALA. First, as a supervised learning model, its recognition ability is confined to the KO families represented in its training data, and it cannot discover new protein families or remote homologs. Second, for the annotation of multi-domain proteins, our proposed Multi-domain mode relies on a hybrid strategy with HMMER, which compromises the tool’s independence. Third, if a given KO is represented by very few protein sequences in the KEGG GENES training data, the model’s detection accuracy for that KO may be reduced. While limitations remain, DeepKOALA demonstrates significant strengths in computational efficiency and achieves a performance comparable to other KO annotation tools. We have uploaded DeepKOALA as an open-source tool on GitHub, and we recommend using the fragment model for cases where sequence length can vary from that in the reference KEGG GENES database, such as metagenomic investigations. Furthermore, while the model is highly optimized for KO annotation, it has been trained on a large and diverse set of labeled protein sequences. This foundational training could allow users to fine-tune the model for other, related downstream classification tasks. Future work is expected to further refine the tool and expand its applicability, including, for this purpose, exploring transfer learning techniques connecting to a new application. DeepKOALA is available on GenomeNet (https://www.genome.jp/tools/deepkoala/) and can be downloaded from our GitHub repository (https://github.com/zhaoxi120/deepkoala).

## Data & Code Availability

The DeepKOALA source code and pretrained model weights are maintained on GitHub (https://github.com/zhaoxi120/deepkoala) and archived at Zenodo (10.5281/zenodo.17949963). This archive also hosts the modified KOfam HMM library (excluding proteins from the 654 held-out species; see Supplementary Table S4) and the dataset of 152 RNA polymerase sequences used for multi-domain evaluation. Due to licensing restrictions, KEGG-derived sequence FASTA files are available upon request for KEGG-subscribed users.

## Author Contribution

Zhaoxi Yu (Conceptualization [lead], Methodology [lead], Data curation [lead], Investigation [lead], Formal analysis [lead], Software [lead], Visualization [lead], Writing—original draft [lead], Writing—review & editing [supporting]), Lingjie Meng (Conceptualization [supporting], Methodology [supporting], Investigation [supporting], Formal analysis [supporting], Writing—original draft [supporting], Writing—review & editing [supporting], Supervision [supporting]), Canh Hao Nguyen (Methodology [supporting], Writing—review & editing [supporting]), Hiroshi Mamitsuka (Methodology [supporting], Writing—review & editing [supporting]), Minoru Kanehisa (Writing—review & editing [supporting], Supervision [supporting], Resources [lead]), Hiroyuki Ogata (Conceptualization [supporting], Methodology [supporting], Formal analysis [supporting], Writing—original draft [supporting], Writing—review & editing [lead], Supervision [lead], Funding acquisition [lead]).

## Funding

This work was supported by JSPS/KAKENHI (No. 22H00384) and a research grant from HFSP (Ref.-No: RGP011/2024; https://doi.org/10.52044/HFSP.RGP0112024.pc.gr.194158).

Computational work was completed at the SuperComputer System, Institute for Chemical Research, Kyoto University.

## Acknowledgements

We thank Koichi Ohkubo and Hideya Uehara for developing, deploying, and maintaining the GenomeNet web interface for the tool described in this manuscript, including ongoing monthly updates aligned with database releases. We also thank Russell Young Neches for helpful suggestions on the experimental design during the early stage of this study.

