## Supplementary Methods for "DeepKOALA: A Fast and Accurate Deep Learning Framework for KEGG Orthology Assignment"

### Supplementary Method

##### Supplementary Method S1 – Data processing and splitting pipeline

###### Data Sources and Versions

- **KEGG GENES database:** February 2025 release.
- **GenomeNet/nr-aa database:** February 2025 release.

###### Command Lines and Parameters

- **DIAMOND (2.1.8):** Command used for homology search to augment sparse KO classes.
  diamond blastp --db nr-aa.dmnd --query K00001.fasta --out diamond_K00001.txt -e 1e-10 --query-cover 80 --subject-cover 80 --id 80 --outfmt 6 -k 100 --threads 4
- **CD-HIT (4.8.1):** Command used for sequence de-redundancy.
  cd-hit -i K00001.fasta -o cdhit_K00001.fasta -c 0.9 -n 5

###### Data Processing Workflow

1. **Initial Data Collection:** A total of 29,207,030 labeled and 26,588,382 unlabeled protein sequences were collected from the KEGG GENES database.
2. **Data Augmentation:** We defined “sparse KO classes” as KOs with fewer than 100 labeled sequences. For each sequence in these sparse KO classes, we performed a DIAMOND (version 2.1.8) search against the GenomeNet nr-aa (non-redundant amino acid sequence) database and added homologous sequences that met strict criteria (E-value < 1e-10, identity ≥ 80%, query and subject coverage ≥ 80%). After augmentation, each sparse KO class was capped at a maximum of 100 labeled sequences.
3. **De-redundancy:** The combined dataset was processed with CD-HIT, resulting in a final non-redundant dataset of 13,682,867 labeled and 18,034,095 unlabeled protein sequences. The sequence length distribution of the full dataset is shown in Supplementary Figure S1.

###### Dataset Splitting Logic

All splits used randomized shuffling and stratification by KO class on the labeled pool. When a split includes unlabeled (“unknown”) sequences, the unlabeled pool is sampled at the same overall percentage that is applied to the labeled pool for that split. This is an equal-percentage sampling rule rather than a fixed 1:1 count.

- **Hyperparameter (HP) phase — 10% subset.** We allocate 7% to HP-train and 3% to HP-validation. HP-train contains labeled (known) proteins only and includes 1,005,589 sequences. It is used exclusively for supervised optimization. HP-validation is mixed under the equal-percentage rule and contains 397,769 labeled sequences and 541,022 unlabeled sequences, for a total of 938,791. There is no ID overlap between HP-validation and any training or evaluation split. Unlabeled sequences in HP-validation are used only to compute open-set criteria such as OpenAUC trade-offs. They never contribute gradients.
- **Five-fold cross-validation (CV) — 90% subset.** Within the remaining 90%, each fold uses 80% train–validation and 20% test. The 80% block is split into 70% train and 10% validation. These sets contain labeled data only. The test set contains the fold’s labeled test portion plus an unknown-test sample drawn from the unlabeled pool using the same percentage as the fold’s known test share. There is no ID overlap with that fold’s train, validation, or known test sets. Per-fold sizes and ratios are summarized in Supplementary Table S1.

###### Oversampling strategy for sparse KO classes in the training sets

After CD-HIT de-redundancy and data augmentation, we applied oversampling to reduce extreme class imbalance in all training sets used across the three training stages (hyperparameter tuning, cross-validation, and final retraining). Oversampling was always applied within the training set only; validation and test sets were left unchanged.

- **Hyperparameter tuning (10% subset).** For this phase, we defined rare KO classes as those with fewer than 7 sequences in HP-train (i.e. 10% of the global target of 70). For each such KO, we randomly resampled its existing sequences with replacement within HP-train until the class contained exactly 7 sequences.
- **Five-fold cross-validation (90% subset).** The main cross-validation experiments used the remaining 90% of the data. In each fold, for the training portion of the fold, KO classes with fewer than 63 sequences (90% of 70) were oversampled in the same way, by randomly duplicating existing sequences until the class contained 63 training sequences.
- **Final retraining (full 70% training set).** For the final model retraining on the full dataset with a 70%/10%/20% train/validation/test split, we defined the target as 70 training sequences per KO class. For any KO class with fewer than 70 sequences in the final training set, we randomly resampled its existing members with replacement until the class contained exactly 70 training sequences.

##### Supplementary Method S2 – Detailed benchmarking environment and setup

###### Software and Hardware Environment

DeepKOALA training, testing, and inference ran on an HPE DL380 with an Intel Xeon Platinum 8462Y+ (Sapphire Rapids) CPU using 8 cores, 150 GB RAM, and one NVIDIA H100 80 GB (PCIe) GPU.

Baseline tool benchmarking on the 60-species test set ran on an HPE Apollo 2000 with an Intel Xeon Gold 6348 (Ice Lake) CPU. DeepKOALA, BlastKOALA, and KofamScan used 32 CPU cores and 128 GB RAM. GhostKOALA used 32 CPU cores and 500 GB RAM.

Software versions and library dependencies used in this study are summarized in Supplementary Table S2.

###### Timing protocol

All local CPU and GPU runs were submitted through a PBS scheduler. For each job, we read the runtime from the PBS completion e-mail using the resources_used.walltime field. This reports on-node elapsed time and excludes queue waiting. Multi-step pipelines report the sum of the steps’ walltimes taken from their respective completion e-mails. Database downloads and one-time index builds are not included. For web-only services (BlastKOALA and GhostKOALA) we report the client-side wall-clock from submission to result retrieval as described below.

###### Baseline Tool Commands and Databases

- **KofamScan:**
  - **Database Version:** February 2025
  - **Command:**
    exec_annotation -o cross_species_comparison.txt cross_species_test.fasta --cpu=32 -f detail-tsv -p modified_profiles -k modified_ko_list
  - KofamScan assignments were accepted only when the best-hit HMM score exceeded the KO-specific adaptive score threshold defined in the ko_list file (modified_ko_list); hits below the threshold were treated as no assignment and were not counted as “recognized”.
- **BlastKOALA and GhostKOALA:**
  - **Database Version:** October 2024
  - **Availability:** These tools were tested using internal KEGG servers; therefore, public command lines are not available.

###### 60-Species Test Set Construction and Database Curation

To ensure a rigorous cross-species evaluation, we curated the reference databases for all baseline tools so that no protein from any species belonging to the same genus as any of the 60 test species remained. In total, 654 species were removed. For KofamScan, this exclusion was applied at the HMM construction stage before multiple alignment and hmmbuild. For BlastKOALA and GhostKOALA, the same species list was removed from the KO reference set on our internal KEGG deployments. No other modifications were made; all remaining steps and parameters followed the official pipelines, and the query sets were unchanged. The complete list of the 60 test species is provided in Supplementary Table S3, and the list of the 654 removed species is provided in Supplementary Table S4.

##### Supplementary Method S3 – Implementation details of fragment model and multi-domain mode

###### Fragment Model (DeepKOALA-fragment)

This model was trained to improve robustness on incomplete sequences. During training, input sequences are processed by the random_cut function. With a 30% probability, the full-length sequence is used. Otherwise (70% probability), a fragment is randomly extracted. The fragment length is chosen from a uniform distribution between a minimum of 50 amino acids and the original sequence length. The Python implementation is as follows:

import random

def random_cut(sequence):
 min_length=50
 full_length_prob=0.3

 sequence = sequence.replace('\n', '')
 seq_len = len(sequence)

 if random.random() < full_length_prob:
 return sequence

 if seq_len < min_length:
 return sequence

 frag_length = random.randint(min_length, seq_len)
 start_pos = random.randint(0, seq_len - frag_length)
 fragment_seq = sequence[start_pos: start_pos + frag_length]

 return fragment_seq

###### Multi-domain mode

This mode uses an iterative hybrid strategy to identify multiple functional domains in a single protein. The workflow is as follows:

1. The full-length sequence (or remaining fragment) is annotated by the DeepKOALA model.
2. If a KO is predicted with a probability above its pre-defined F1-optimized threshold, the corresponding HMM profile from KofamScan is used with HMMER to identify the precise domain boundaries.
3. The identified domain is computationally excised, and any remaining fragments longer than 50 amino acids are added to a queue for re-analysis.
4. The process repeats, taking the next fragment from the queue and feeding it back to step 1, until no fragments longer than 50 amino acids remain.

**HMMER Search Parameters:** The hmmsearch command is run with default reporting thresholds (i.e., no specific E-value or bit score cutoffs are applied via command-line flags). The algorithm parses the domain table output (--domtblout) and extracts the boundaries of the top-ranked hit, which corresponds to the hit with the lowest E-value. This strategy is based on the premise that the initial DeepKOALA prediction provides sufficient confidence to guide the boundary search.

**Version Consistency:** For optimal performance and reproducibility, the HMM profiles used by Multi-domain mode should correspond to the same database release that the DeepKOALA model was trained on. The models and results presented in this study are based on the February 2025 versions of both the DeepKOALA model and the KofamScan HMM profiles.

##### Supplementary Method S4 – Detailed per-KO threshold-setting pipeline

The optimal decision threshold for each KO class was determined as follows:

1. The trained model was used to generate predictions and class-specific Softmax scores on the entire training and validation sets.
2. For each KO class, we focused on the sequences whose predicted label was that class and divided them into two groups: sequences whose true label matched the KO class, and sequences whose true label differed (including sequences annotated with other KO classes or unlabeled sequences).
3. Using the class-specific Softmax score distributions of these two groups, we swept over all distinct Softmax scores as candidate thresholds, computed precision and recall for that KO class at each threshold, and selected the score that maximized the F1-score. For KO classes with degenerate prediction patterns, we applied the following fallback rules when computing thresholds. First, if there were predicted sequences for a KO class but none of them had that KO as the true label (i.e. the “correct” group was empty and only the “incorrect” group was present), we set the threshold to the maximum Softmax score observed among the incorrectly predicted sequences. Second, if there were only correctly predicted sequences for a KO class (i.e. the “incorrect” group was empty), we set the threshold to the minimum Softmax score observed among the correctly predicted sequences. Finally, if the model never predicted a given KO class on the training and validation sets, we assigned that class a default global threshold estimated from the pooled Softmax scores across all KO classes.

The relationship between per-KO test F1-scores and training sample size is illustrated in Supplementary Figure S2.

##### Supplementary Method S5 – Extended metrics and hyperparameter optimization

**Hyperparameter Optimization Protocol**

We conducted model-specific hyperparameter optimization prior to cross-validation. For the RNN, GRU, and LSTM models, the search spaces are summarized in Supplementary Table S5 (classical RNN family block), and we executed 15 trials per model. During preliminary screening we observed that GRU with hidden_size = 192 reproducibly triggered CUDA runtime errors on our GPU setup; therefore, this single value was excluded from the GRU search space. For the Mamba architecture, the search space is listed in Supplementary Table S6 (Mamba block). We attempted 30 trials, of which 23 completed successfully (the remainder terminated due to runtime failures and are not included in the results). For ESM-2 (8M), we only tuned the classifier MLP that follows the frozen backbone; the learning rate was selected from {1e-3, 5e-4, 1e-4}.

Across all models, we used Bayesian optimization with OpenAUC as the objective, operating on the HP-train/HP-validation split described in Supplementary Method S1. We set the maximum epochs to 50 with early stopping (min_delta = 1e-3, patience = 5) and applied gradient clipping with a maximum norm of 5. Unless otherwise specified in Supplementary Table S5 and S6, other training hyperparameters followed the defaults of our training framework. The complete trial-wise results for each model are reported in Supplementary Table S7, and fold-wise metrics of the selected configurations are provided in Supplementary Table S8.
