## Supplementary Figures for "DeepKOALA: A Fast and Accurate Deep Learning Framework for KEGG Orthology Assignment"

### Supplementary Figure

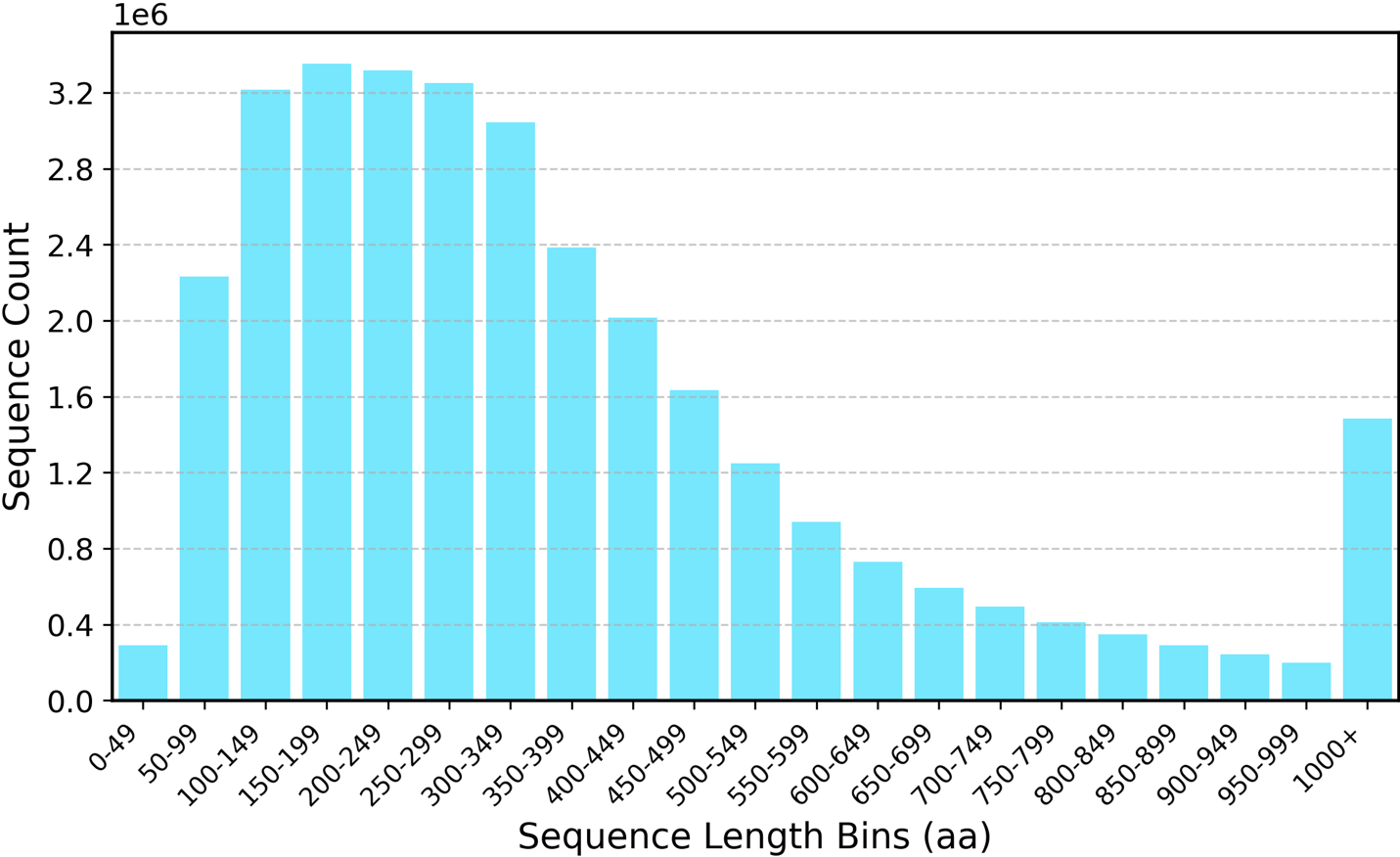

**Supplementary Figure S1** – **Sequence Length Distribution of the Full Dataset**

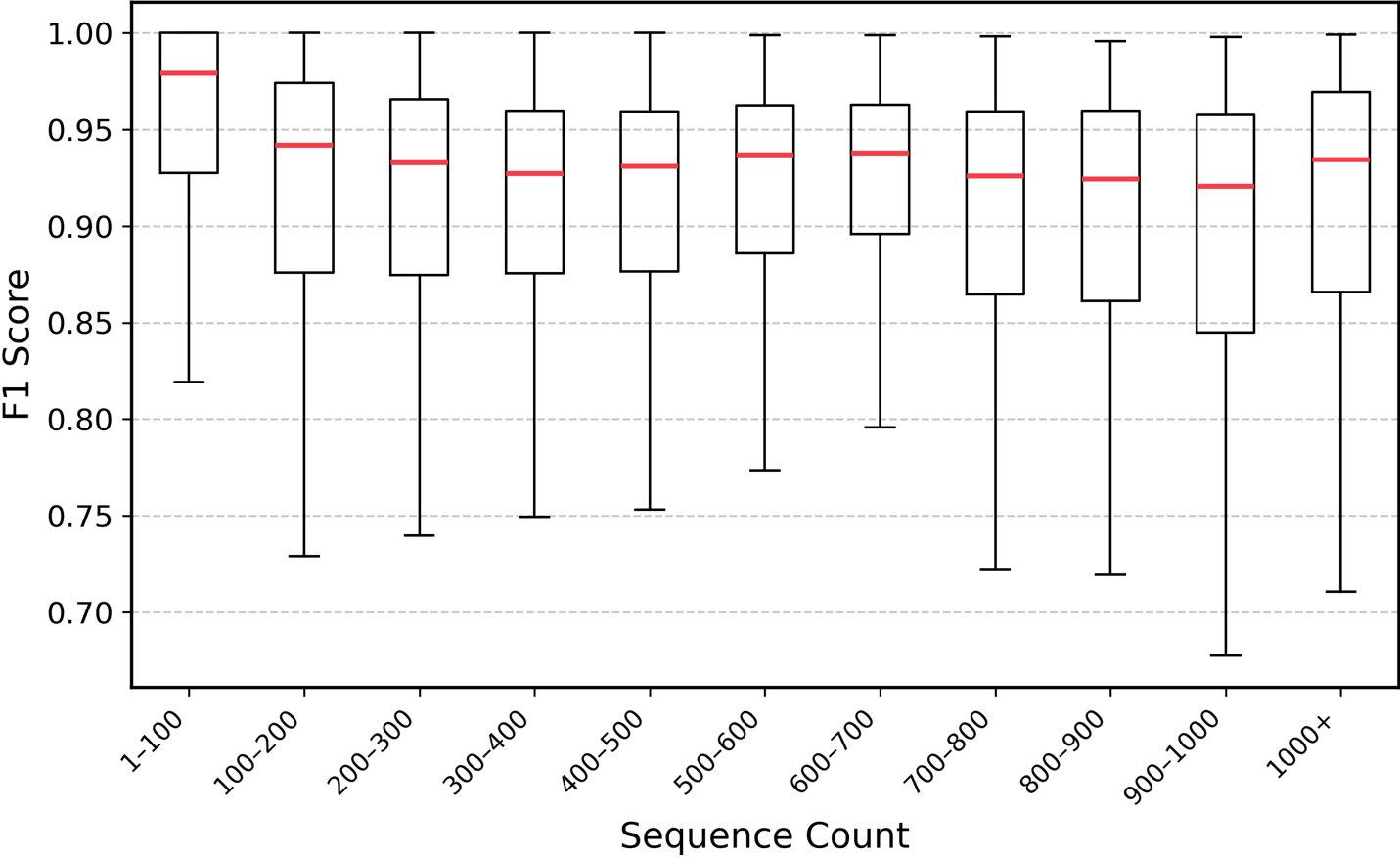

**Supplementary Figure S2** – **Per-KO test F1 versus sample size**

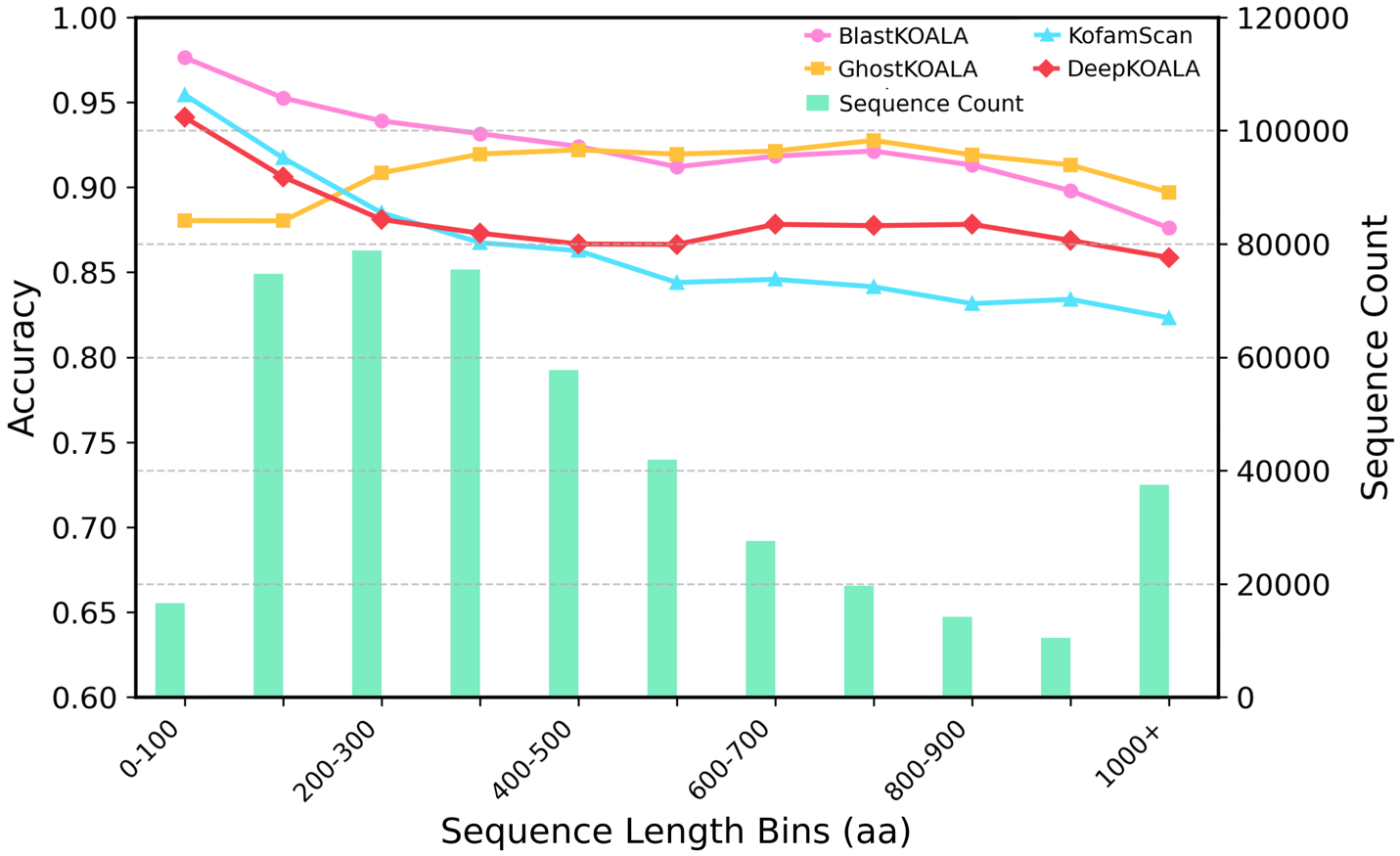

**Supplementary Figure S3** – **Accuracy versus sequence length on the 60-species test set**
