## Supplementary Tables for "DeepKOALA: A Fast and Accurate Deep Learning Framework for KEGG Orthology Assignment"

### Supplementary Table

**Supplementary Table S1: Per-fold sample counts for five-fold cross-validation (train/validation/test)**

|  | Train | Validate | Test |
| --- | --- | --- | --- |
| Fold1 | 9,111,427 | 1,255,471 | 5,748,310 |
| Fold2 | 9,113,137 | 1,257,999 | 5,743,198 |
| Fold3 | 9,116,587 | 1,257,999 | 5,738,258 |
| Fold4 | 9,120,147 | 1,257,999 | 5,733,165 |
| Fold5 | 9,123,630 | 1,257,999 | 5,728,354 |

**Supplementary Table S2** – **Software and Library Versions**

| **Software Name** | **Version** | **URL** |
| --- | --- | --- |
| Python | 3.11.9 | https://www.python.org/ |
| PyTorch | 2.4.1 | https://pytorch.org/ |
| Pandas | 2.2.2 | https://pandas.pydata.org/ |
| Numpy | 1.26.3 | https://numpy.org/ |
| Scikit-learn | 1.2.2 | https://scikit-learn.org/stable/ |
| DIAMOND | 2.1.8 | https://github.com/bbuchfink/diamond |
| CD-HIT | 4.8.1 | https://sites.google.com/view/cd-hit/home |
| KofamScan | 1.3.0 | https://github.com/takaram/kofam_scan |
| HMMER | 3.4 | http://hmmer.org/ |
| CUDA | 12.7 | https://developer.nvidia.com/cuda-toolkit |

**Supplementary Table S3** – **List of 60 species in the held-out test set**

**Supplementary Table S4 — Species removed from train/validation due to genus-level overlap with the test set (n = 654)**

**Supplementary Table S5 – Hyperparameter search spaces for RNN, GRU, and LSTM**

|  | Hidden size | Layer number | Learning rate |
| --- | --- | --- | --- |
| RNN | 64, 96, 128, 192, 256 | 1, 2 | 1e-3, 5e-4, 1e-4 |
| GRU | 64, 96, 128, 256 | 1, 2 | 1e-3, 5e-4, 1e-4 |
| LSTM | 64, 96, 128, 192, 256 | 1, 2 | 1e-3, 5e-4, 1e-4 |

**Supplementary Table S6** – **Hyperparameter search space for Mamba**

|  | D_model | D_state | D_conv | expand | Learning rate |
| --- | --- | --- | --- | --- | --- |
| Mamba | 64, 96, 128, 192, 256 | 8, 16 | 3, 4 | 1, 2, 3 | 1e-3, 5e-4, 1e-4 |

**Supplementary Table S7 – Complete hyperparameter optimization results (trial-wise) across models**

**Supplementary Table S8 — Five-fold cross-validation performance of the selected configurations**

**Supplementary Table S9 — Detailed DeepKOALA and KofamScan annotations for the 152 RNA polymerase test set**
